# Quantifying spatial dynamics and regulators of *Mycobacterium tuberculosis* phagocytosis by primary human macrophages using microfabricated patterns

**DOI:** 10.1101/2022.11.10.515919

**Authors:** Anca F. Savulescu, Nashied Peton, Delia Oosthuizen, Rudranil Hazra, Musa. M. Mhlanga, Anna K. Coussens

## Abstract

Macrophages provide a first line of defense against invading pathogens, including the leading cause of bacterial mortality, *Mycobacterium tuberculosis* (*Mtb*). Phagocytosing extracellular organisms mediate pathogen clearance *via* a multitude of antimicrobial mechanisms, uniquely designed against an array of pathogens. Macrophages are able to execute different programs of activation in response to pathogenic challenge with host mediators, polarizing them to a variety of differentiation states, including the pro-inflammatory M1 and anti-inflammatory M2 states. The functional polarization of a macrophage prior to infection, thus impacts the outcome of host-pathogen interaction. One of the limitations when using *in vitro* differentiated human primary monocyte-derived macrophages (MDMs) is the heterogeneous nature of the mature population, which presents a challenge for quantitative characterization of various host-pathogen processes. Here, we describe an approach to minimize this heterogeneity, based on micropatterning of cells to reintroduce aspects of cellular homogeneity lost in a 2D tissue culture. Micropatterning consists of growing cells at the single cell level on microfabricated patterns, to constrain the size and shape of the cell, reducing cell-to-cell variation and mimicking the physiological spatial confinement that cells experience in tissues. We infected micropatterned GM-CSF- (M1) and M-CSF- (M2) derived human MDMs with *Mtb*, which allowed us to study host-pathogen interactions at a single cell level, at high resolution and in a quantitative manner, across tens to hundreds of cells in parallel. Using our approach, we were able to quantify phagocytosis of *Mtb* in MDMs, finding phagocytic contraction is increased by interferon-gamma stimulation, whilst contraction and bacterial uptake is decreased following silencing of phagocytosis regulator *NHLRC2* or Tween80 removal of bacterial surface lipids. We also identify alterations in host organelle position within *Mtb* infected MDMs, as well as identifying differences in *Mtb* subcellular localization in relation to the microtubule organizing center (MTOC) and in line with the cellular polarity in M1 and M2 MDMs. Our approach described here can be adapted to study other host-pathogen interactions and co-infections in MDMs and can be coupled with downstream automated analytical approaches.

## INTRODUCTION

*Mycobacterium tuberculosis* (*Mtb*), the causative agent of tuberculosis disease remains the leading bacterial cause of death world-wide and was responsible for about 1.5 million deaths in 2020 (WHO Global TB report, 2021). Macrophages represent an essential cellular component of the innate immune system responding to both acute and chronic inflammatory processes. These immune cells are the first line of defense for the control of pathogens, limiting spread through activation of surface markers, production of proinflammatory cytokines and chemokines and recruitment of peripheral lymphocytes and monocytes to the site of inflammation. With macrophages being the predominant host cells implicated in bacterial entry, growth and restriction, they have become the immune cell of choice for the study of host-pathogen interactions (Gordon *et al*., 2017). Alveolar macrophages are the primary immune cells to phagocytose inhaled *Mtb* when it reaches the distal airways and thus their functional response is critical to the outcome of infection (Marakalala *et al*., 2018).

There are two major lineages of macrophage polarization, pro-inflammatory M1 (classically activated) and anti-inflammatory M2 (alternately activated) phenotypes, although additional M2 subphenotypes and other lineages such as M3 and M4 have also been proposed (de Sousa *et al*., 2019). Structurally, M1 form a more round cellular phenotype, whilst M2 are more filamentous and heterogenous in shape (McWhorter *et al*., 2013, Hu *et al*., 2021). Heterogeneity in macrophages induced by mycobacterial infection is marked by altered expression of pro-inflammatory or anti-inflammatory responses (Verreck *et al*., 2004). This heterogeneity is particularly relevant when studying complex host-pathogen interactions, when not all cells in a tissue culture dish will be infected, nor undergoing the same process at the same time. Although most studies have routinely used *in vitro*-generated human monocyte-derived macrophage (MDMs) with great success to analyze cellular responses at a bulk population level, their capacity to differentiate into a homogenous population remains elusive due to their high degree of plasticity and ability to assume different phenotypes depending on the local microenvironment (Gordon *et al*., 2005, Martinez *et al*., 2008, Gordon *et al*., 2011). Quantification at the single cell level mitigates some of the limitations of bulk analyses, however, random dispersal of heterogeneous cells across a dish still makes systematic quantification of individual cell responses problematic.

To overcome these difficulties we made use of micropatterning of cells, which imposes morphological homogeneity in cell shape and size, and spatial organization of organelles (Thery *et al*., 2005, Thery *et al*., 2006^a^, Schauer *et al*., 2010., Savulescu *et al*., 2020), mimicking the physiological spatial confinement that cells experience in tissues. Micropatterning leads to reduction in cell-to-cell variability and allows the averaging of a high number of cells. This allows quantitative study of biological processes, including spatial and temporal subcellular localization of biomolecules (Savulescu *et al*., 2021). We made use of this system to study early events during *Mtb* infection of GM-CSF-(M1) and M-CSF-(M2) derived human MDMs at a high resolution, in a quantitative manner and on a single cell level.

*In vitro* 1D micropatterned adhesive substrates and adhesive microfabricated patterns have been previously used to grow mouse bone marrow-derived and peritoneal macrophages (Sharma *et al*., 2012, Sharma *et al*., 2014, Trescos *et al*., 2015). Trescos *et al*., treated bone marrow-derived macrophages (BMDMs) and primary peritoneal macrophages grown on micropatterns, with the edema toxin (ET) - the major virulence factor of the Anthrax causing bacterium - *Bacillus anthracis*. ET induced a dynamic disruption of the actin cytoskeleton, eventually leading to cell shrinkage and complete disruption of the cell architecture, including relocalization of the nucleus (Trescos *et al*., 2015). Micropatterns have also been used to analyze F-actin cytoskeleton restructuring induced by infection of the HeLa cell line with *Salmonella Typhimurium* (*Stm*) demonstrating that docking and invasion sites of *Stm* were preferred to specific cellular height (Vonaesch *et al*., 2013). In these studies mentioned above, micropatterning of cells allowed for standardization and quantification of the biological process studied, including cytoskeleton rearrangements and bacterial infection, as well as statistical cell-to-cell comparison among the cellular population.

Here, we introduce a novel high-throughput, single-cell and microscopy-based method to study early stages of *Mtb* infection in primary human MDMs, based on micropatterning of cells. We seed and grow MDMs on two types of microfabricated patterns - standard and elastomeric, followed by infection with *Mtb* and fixation at various time points. Standard patterns allow quantification of the subcellular location of *Mtb* bacilli relative to organelle location; elastomeric micropatterns have the ability to contract in reaction to a mechanical force applied onto the adhered cell, allowing quantification of cellular contraction during phagocytosis. We quantified *Mtb* phagocytosis and assessed the impact of culturing in an interferon-gamma (IFN-γ) pro-inflammatory microenvironment, silencing macrophage expression of *NHLRC2*, a key regulator of actin polymerisation, and striping bacterial cell wall lipids, on phagocytosis. Using standard micropatterns, we show that *Mtb* infection alters the position of the microtubule organizing center (MTOC) in opposite manners in M1 and M2 MDMs, as well as altering the position of the cell nucleus in both MDM types. Finally, we demonstrate a correlation between the subcellular location of *Mtb* bacilli and the MTOC-positioning, as indicative of cellular polarity. In summary, we were able to visualize and quantify these processes due to the homogenous nature of the cell populations once seeded on the micropatterns. Our method can be readily applied to the high throughput image-based study of additional pathogen infections in MDMs, allowing for quantitative studies of spatial and temporal subcellular localization of host-pathogen interactions.

## RESULTS

### *Mtb* H37Rv infection of MDMs on micropatterns

We first set out to establish a micropatterned-based assay to reduce variation in cell-to-cell structural and organelle spatial organization, which would allow quantitative study of early stages of *Mtb* infection in MDMs. To do so, we isolated CD14+ monocytes from healthy donor PBMCs and then differentiated them *in vitro* for 7 days in the presence of either GM-CSF or M-CSF to generate M1 and M2 MDM, respectively (Figure 1A). At the site of *Mtb* infection, macrophages are exposed to a cytokine-enriched microenvironment, with interferon-gamma (IFN-γ) produced by *Mtb*-specific T cells, a hallmark of TB inflammation (Kumar, 2017). MDM were therefore cultured in the presence or absence of IFN-γ before infection. We then made use of the FLECS (fluorescently labeled elastomeric contractible surfaces) technology (Pushkarsky *et al*., 2018) to measure phagocytosis of *Mtb* in MDMs (Figure 1A). In this technique, cells are grown on elastomeric micropatterns and mechanical forces, such as mechanotransduction, phagocytosis and others, are measured as contractions in the elastomeric micropatterns and quantified at a single cell level. We seeded MDMs on fluorescently-labeled elastomeric micropatterns at a concentration of 75,000 cells/0.5 ml per well in 24-well plates. The cells were left to adhere for 60-90 minutes, followed by removal of non-adhesive cells by gentle washing. The macrophages were then infected with mCherry fluorescently labeled *Mtb* at a multiplicity of infection (MOI) of 10 for 4 hours. This was followed by fixation of cells overnight at 37°C with 4 % pre-warmed paraformaldehyde (PFA), 1x PBS washes and storage at 4°C. Once fixed, we performed immunofluorescence to label actin and DNA, and imaged the cells on a StellarVision microscope, using Synthetic Aperture Optics (SAO). The length of each elastomeric pattern was measured manually using FiJi. Only single cells that spread out appropriately on the micropattern were quantified.

**Figure 1.**
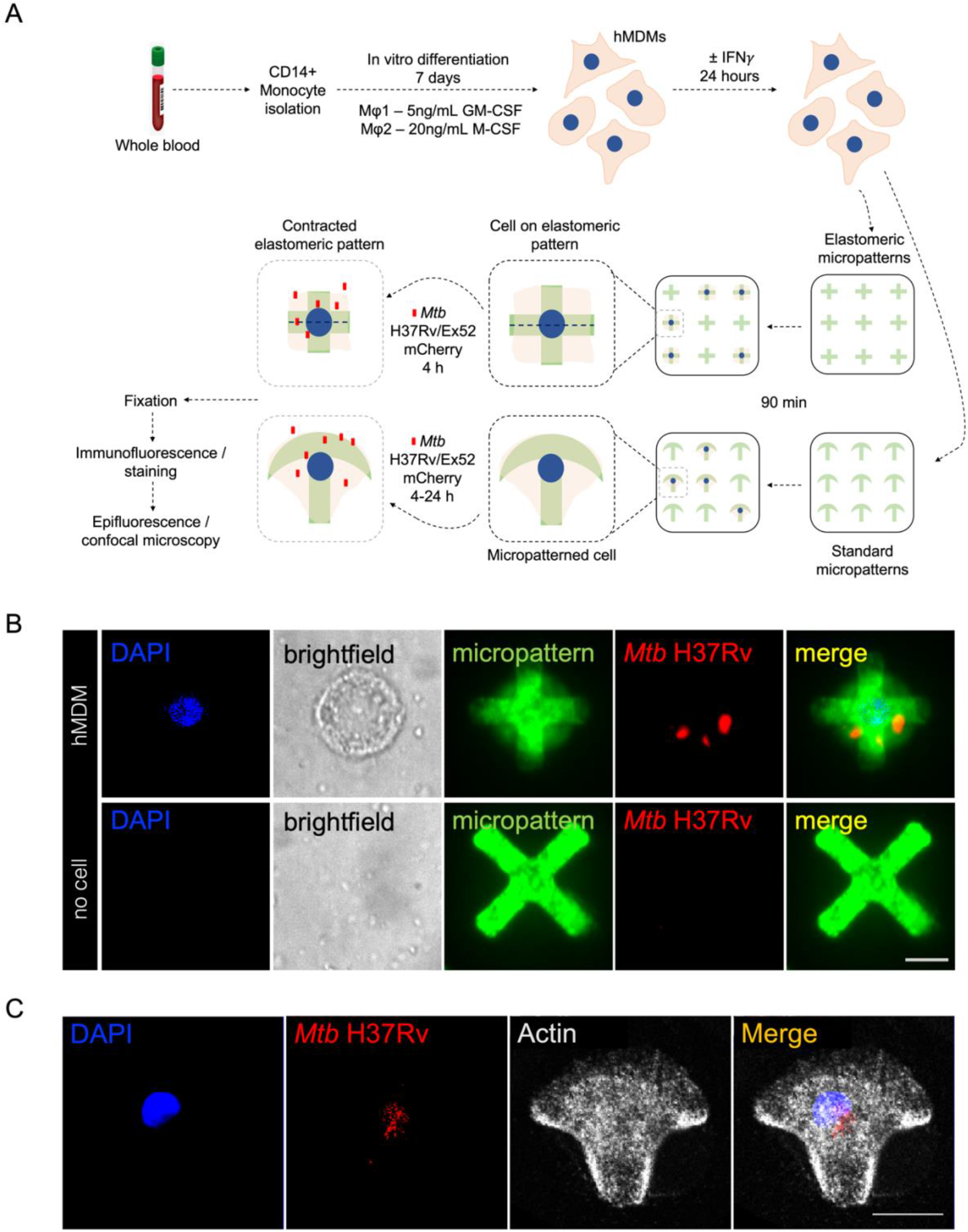
Methodology schematic for *Mtb* infection of human MDMs using elastomeric and standard micropatterns. **A**. PBMCs are isolated from whole blood of healthy donors by gradient separation followed by CD14+ monocyte isolation from PBMC using magnetic bead positive selection. CD14+ monocytes are differentiated into M1 and M2 macrophages followed by overnight stimulation with or without 100 IU/mL IFN-γ. Cells are then seeded at a concentration of ∼75,000 cells per 0.5 ml onto elastomeric or standard patterns, left to adhere and adopt the appropriate shape for 60-90 minutes, with unbound cells washed prior to infection. Cells are infected with either *Mtb* laboratory strain H37Rv, or clinical strain EX52 for 4 hours, and cells washed and either fixed immediately, or cultured for up to 24 hours. Cells are then fixed for 16-24 hours overnight, washed with 1x PBS and taken for immunofluorescence, followed by imaging using epifluorescence/confocal microscopy. **B**. Images of a *Mtb* H37Rv infected M1 MDM on an elastomeric micropattern (top) and an elastomeric pattern that does not contain a cell (bottom). Contraction of the elastomeric pattern is clearly seen in the top, but not in the bottom image. DNA is stained with DAPI (blue), the cell is shown in brightfield (gray), the micropattern is labeled with Alexa Fluor 488 (green) and *Mtb* H37Rv expresses mCherry (red). A merged image is shown on the right. Scale bar 10 μm. **C**. An image of a *Mtb* H37Rv infected M1 MDM on a standard crossbow micropattern. DNA is stained with DAPI (blue), *Mtb* H37Rv expresses mCherry (red), and actin is stained with phalloidin (gray). A merged image is shown on the right. Scale bar 10 μm.

In parallel, we prepared micropatterns based on the protocol described in Azioune *et al*., 2009. Briefly, we coated coverslips with poly(L-lysine) poly(ethylene glycol) (PLL-g-PEG) and made use of a chromium photomask to imprint the desired size and shape of micropatterns onto coated 10 mm coverslips. We then coated the imprinted micropatterns with fibronectin, followed by seeding of MDMs at a concentration of 75,000 cells/0.5 ml per coverslip in 24-well plates. Micropattern sizes ranging from small to large were tested, as well as two different shapes (crossbow and round) (Figure S1). However, medium-sized crossbow shaped micropatterns were our preferred pattern of choice as they have been shown to promote a polar intracellular organization of cells (Thery *et al*., 2006^I^, Schauer *et al*., 2010). Additionally their size is typical to that of MDMs. The cells were left to adhere for 60-90 minutes, followed by removal of non-adhesive cells by gentle washing. Both cell types adhered well to the micropatterns, but we found that pre-treatment with 100 IU/ml IFN-γ for 24 hr prior to seeding, improved cellular spreading and adoption of the correct micropatterned shape, for both cell types. Following removal of the non-adherent cells, macrophages were infected with mCherry/GFP fluorescently labeled *Mtb* at a multiplicity of infection (MOI) of 10 for 4 hours. This was followed by fixation of cells at 4, 16, or 24 hours post infection using 4 % pre-warmed PFA, 1x PBS washes and storage at 4°C. We then performed immunofluorescence and epifluorescent or confocal microscopy on fixed cells to visualize *Mtb* bacilli and cellular landmarks, including the MTOC and actin structure. This was followed by manual image analysis using the FiJi program to characterize the subcellular localization of *Mtb* bacilli in relation to cellular markers. Only cells that adopted the micropattern shape correctly were quantified. We note that the imaging data can be coupled with automated analytical approaches to quantify subcellular localization patterns and colocalization levels, such as in Savulescu *et al*., 2021. Images of typical MDMs on elastomeric micropatterns and standard crossbow micropatterns are shown in Figure 1B-C.

### Elastomeric micropatterns provide a quantitative assay to measure *Mtb* H37Rv phagocytosis in MDMs

We made use of the FLECS technology to specifically look at the earliest stage in *Mtb* infection, namely phagocytosis. The elastomeric patterns are cross-shaped, in a 24-well plate format, coated with fibronectin and Fibrinogen-Alexa Fluor 488 fluorescent dye. Upon exertion of a mechanical force on cells growing on these micropatterns, the cells and subsequently, the elastomeric micropatterns contract. We seeded MDMs on elastomeric patterns, allowed them to adhere and adopt the typical shape for 90 minutes, washed off unadhered cells and infected them with *Mtb* H37Rv (a common lineage 4 laboratory strain of *Mtb)* expressing mCherry, for 4 hours, followed by fixation. We then stained fixed samples with DAPI and imaged the cells on a StellarVision microscope. The length of the elastomeric patterns was measured manually, using FiJi. Initially, we set out to test whether phagocytosis of H37Rv can be visualized and quantified using these elastomeric patterns. We compared IFN-γ stimulated cells with unstimulated cells, as well as H37Rv infected cells with uninfected cells and combinations of these conditions (Figure 2A-B). Unstimulated cells that were infected with H37Rv showed a higher level of contraction compared to unstimulated and uninfected cells (P<0.0001), suggesting that the assay is able to capture phagocytosis. Interestingly, stimulation of cells with IFN-γ in the absence of infection also resulted in contraction, compared to untreated cells, although less significant (P=0.0057). As such, the largest contraction was observed in cells that were both stimulated with IFN-γ and infected with H37Rv (P<0.0001). This suggests that at *in vivo* sites of disease an IFN-γ microenvironment may increase macrophage *Mtb* phagocytosis.

**Figure 2.**
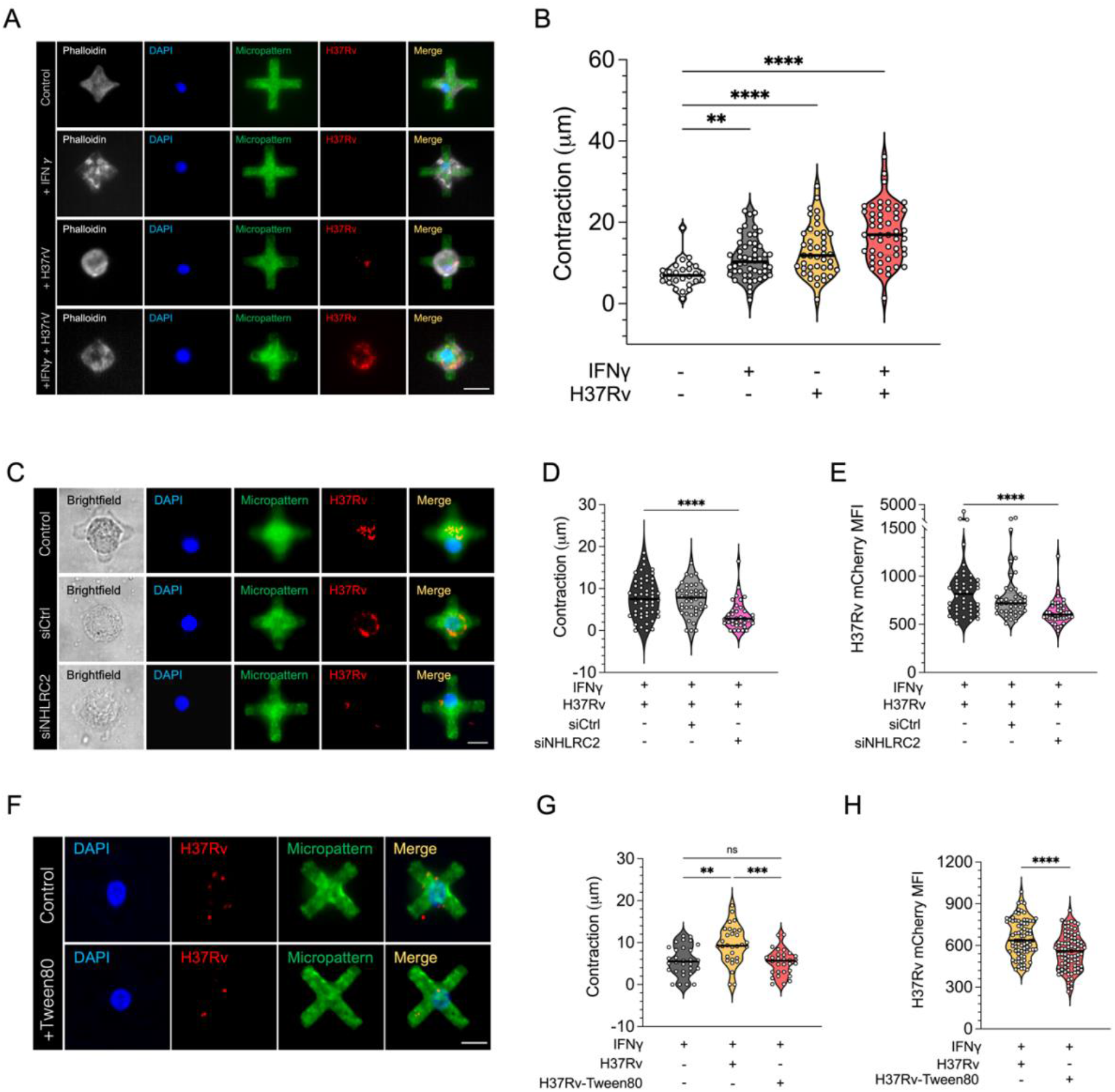
Elastomeric patterns provide a quantitative assay to measure phagocytosis of *Mtb* H37Rv by M1 MDMs. **A**. Images of MDMs unstimulated / stimulated with IFN-γ and uninfected / infected with H37Rv. Actin is stained with Phalloidin (gray), the DNA is labeled with DAPI (blue), *Mtb* bacilli expresses mCherry (red) and the elastomeric pattern is labeled with AlexaFluor-488 (green). A merged image is shown on the right. Scale bar 25 μm. **B**. Contraction (measured in μm) of elastomeric micropatterns in MDMs unstimulated / stimulated with IFN-γ and uninfected / infected with H37Rv. **C**. Images of MDM cell infected with H37Rv (Control), MDM cell infected with H37Rv and transfected with non-specific scramble siRNA (siCtrl) and MDM cells infected with H37Rv and transfected with siNHLRC2 (siNHLRC2). A brightfield image is shown on the left (gray), the DNA is labeled with DAPI (blue), *Mtb* bacilli expresses mCherry (red) and the elastomeric pattern is labeled with AlexaFluor-488 (green). A merged image is shown on the right. Scale bar 20 μm. **D**. Contraction of elastomeric micropatterns in infected MDMs, treated with IFN-γ, that were either transfected with non-specific scramble siRNA (siCtrl), siNHLRC2 or untransfected, n=3 donors. **E**. Mean fluorescence intensity (MFI) of H37Rv in MDM cells, treated with IFN-γ, that were either transfected with siCtrl, siNHLRC2 or untransfected, n=3 donors. **F**. Images of MDMs infected with H37Rv (top) and MDMs infected with H37Rv-Tween80 (bottom). Scale bar 20 μm. **G**. Contraction of elastomeric micropatterns in uninfected MDMs, MDMs infected with H37Rv and MDMs infected with H37Rv-Tween80, n=2 donors. The DNA is labeled with DAPI (blue), *Mtb* bacilli expresses mCherry (red) and the elastomeric pattern is labeled with AlexaFluor-488 (green). A merged image is shown on the right. **H**. MFI of H37Rv or *Mtb* H37Rv-Tween80 in MDM cells, n=2 donors. Violin plot line, median, dotted lines, IQR; analyzed by Kruskall-Wallis test with Dunn’s multiple comparison or Mann Whitney test; *, P<0.05; **, P<0.01; ***, P<0.001; ****, P<0.0001; ns, not significant.

We next sought to validate the ability of our assay to be used for functional assessment of host and pathogen mediated mechanisms governing phagocytosis. To do so, we first used siRNA to deplete expression of *NHLRC2* (siNHLRC2), the top hit in a genome-wide CRISPR screen identified as a key regulator of phagocytosis in the human monocytic cell line U937 (Haney *et al*., 2018). We tested the impact of *NHLRC2* silencing on contraction of IFN-γ-treated-H37Rv-infected MDMs, the condition with the greatest contraction. siNHLRC2 cells showed a significant decrease in contraction (P<0.0001) compared to both cells transfected with non-specific scramble siRNA (siCtrl) and untransfected controls; there was no significant difference between siCtrl and untransfected controls (Figure 2C-D). To confirm that the contraction is indeed a measurement of phagocytosis, we measured the *Mtb* mCherry mean fluorescence intensity (MFI) in siNHLRC2, siCtrl and untransfected cells. In line with the level of contraction, a significantly lower mCherry MFI was observed in siNHLRC2 cells compared to siCtrl and untransfected cells (Figure 2E). Taken together, these data demonstrate the functional validity of our assay to study macrophage mechanism of phagocytosis and confirm a novel role for NHLRC2 as a regulator of *Mtb* phagocytosis by primary human macrophages.

As it has been shown that *Mtb* strain variation in cell wall surface lipids influences receptor-mediated uptake by macrophages (Schlesinger *et al*., 1996), next, we used the elastomeric patterns to test whether stripping lipids off the surface of *Mtb* H37Rv modulates macrophage contraction during phagocytosis. To do so, we cultured H37Rv mCherry in the absence (H37Rv) or presence of Tween80 (H37Rv-Tween80), a detergent commonly used to remove the surface lipids from *Mtb* during broth culture to prevent bacilli from clumping. Single cell suspension stocks for infection were then prepared in an identical manner using glass beads to dissociate bacterial clumps and create a single cell suspension with sequential low-speed centrifugation. Bacterial counts were confirmed by CFU plating and then stocks used to infect MDM at an MOI of 10:1 for 4 hours. Infecting IFN-γ-stimulated M1 MDMs on elastomeric micropatterns with the two preparations of *Mtb*, consistent with Figure 2A, we again saw a significant increase in contraction when infecting with H37Rv grown in the absence of Tween80, compared to IFN-γ-stimulation only (P=0.0016). However, this H37Rv-induced contraction was significantly reduced when infecting with H37Rv-Tween80 (P=0.0004), to a level of contraction similar to IFN-γ-stimulation only (Figure 2F-G). To confirm that the significant difference in contraction observed between the two bacterial preparations is indeed a measurement of phagocytosis, we measured the mCherry MFI. In line with the level of contraction, a higher MFI was observed in cells infected with cell wall lipid intact H37Rv, compared to cells infected with H37Rv-Tween80 (P=0.040) (Figure 2H). This data confirms the utility of our assay to investigate bacterial factors which modify *Mtb* phagocytosis in a quantitative manner, at a single cell level.

### Micropatterns are suitable for the study of early, but not late stages of *Mtb* H37Rv infection in M1 and M2 MDMs

Having used elastomeric patterns to quantify processes involved in Mtb phagocytosis, we next made use of standard micropatterns to quantify various intracellular processes following *Mtb* uptake. First, we set out to characterize the time frame over which we could successfully capture micropatterned MDMs following infection with *Mtb* H37Rv. Typically, studies using micropatterned cells explore processes that occur within a short time frame from seeding (Thery *et al*., 2006^b^, Schauer *et al*., 2010). We therefore wanted to establish this timeframe for MDMs infected with *Mtb*. We seeded M1 and M2 MDMs on crossbow micropatterns and infected them with mCherry/GFP fluorescently labeled H37Rv for various time points ranging from 4 to 24 hours, before fixing. We then stained cells with DAPI and imaged them on a Stellarvision microscope at a low magnification to allow the visualization and quantification of hundreds of cells. We manually counted the number of cells on each coverslip and compared between the numbers at different time points. The typical number of MDMs that were adhered and spread appropriately on micropatterns was ∼ 200-250 cells per 10 mm coverslip (Figure S2). For comparison, we seeded M1s and M2s on both crossbow micropatterns without infecting them, to test whether infection with *Mtb* H37Rv modifies adherence on micropatterns. Typical images of H37Rv infected M1 and M2 cells on crossbow shaped micropatterns are shown in Figure 3A. The DNA is stained with DAPI, the cell is seen in brightfield, the micropatterns are labeled with AlexaFluor-488 and we used an H37Rv fluorescent reporter strain which constitutively expressed mCherry. We set the number of cells that were adhered and appropriately spread on micropatterns at the earliest time point (4 h) as the 100 % and calculated the number of cells at each following time point as a percentage of this 4 h time point. Absolute number of adhered and appropriately spread cells on crossbow micropatterns are shown in Figure S2. No significant differences were observed in the number of adhered and appropriately spread cells at 4 h between uninfected M1s and M2s and between infected M1s and M2s (P=0.832 and 0.476, respectively) (Figure S2). Similarly, no significant differences were observed in adherence and spreading at 4 h between uninfected versus infected M1s or M2s (P=0.725 and 0.772, respectively) (Figure S2). For both M1s and M2s, a declining trend can be seen from 4 h throughout the timeline, to a low of ∼5-15 % cells remaining at the 24 h time point, irrespective of H37Rv infection (Figure 3B). Notably, less uninfected M1s remained adhered to micropatterns compared to infected cells; only ∼55-60 % of uninfected M1s remained adhered to micropatterns at 8 h, compared to 70-80 % for infected M1s. Similarly, a slightly higher number of H37Rv infected M1s remained adhered at the 16 h and 24 h time points, compared to uninfected M1s (Figure 3B). This suggests that infection on M1s with H37Rv may help adhere or/and stabilize these cells on micropatterns. Curiously, the opposite can be seen for uninfected versus H37Rv infected M2s at the 8 h time point - infection with *Mtb* H37Rv reduced the number of adhered cells on micropatterns (Figure 3B). However, this reduction is not consistent at later time points (16 h and 24 h) (Figure 3B). When translating the percentages into cell numbers, the typical number of adhered and properly spread uninfected and H37Rv-infected M1 and M2 MDMs at 8 h was ∼120-130 (Figure S2), which we considered as an appropriate number of cells that allowed quantification and statistical analyses. However, the number of cells that remained adhered on micropatterns at the 16 h and 24 h time points was lower than 75 cells in most cases, which we considered to not be sufficient for quantitative and statistical downstream analyses, as previously shown (Thery *et al*., 2006^a^, Schauer *et al*., 2010, Vonaesch *et al*., 2013).

**Figure 3.**
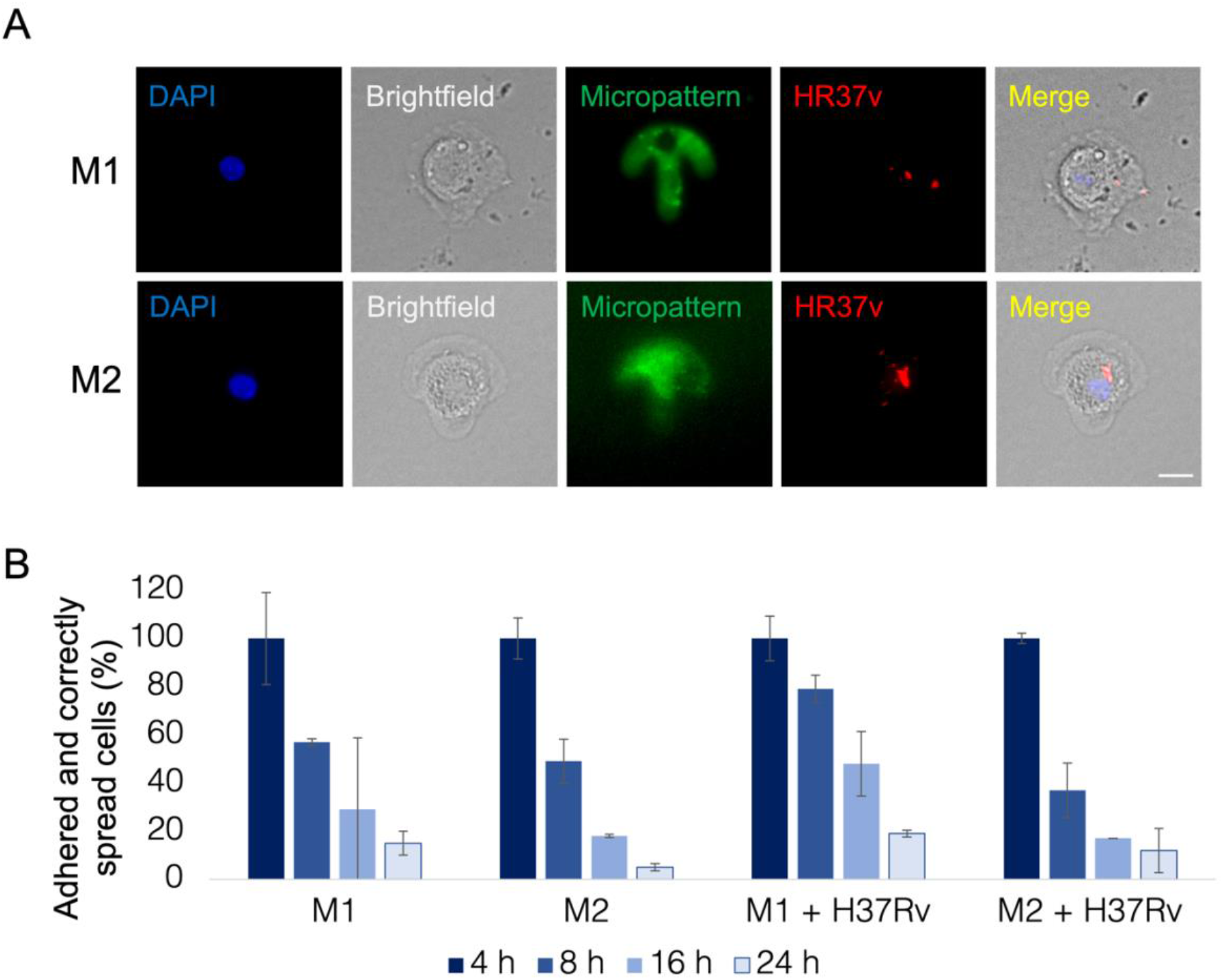
Micropatterns are suitable for early, but not late stages of H37Rv infection in MDMs. **A**. Typical H37Rv infected M1 and M2 MDMs on crossbow shaped micropatterns. The DNA is labeled with DAPI (blue), the cell can be seen in brightfield, micropatterns are labeled with AlexaFluor-488 (green), *Mtb* H37Rv expresses mCherry (red) and a merged image (combining all fields other than the green micropattern, for clarity) is shown on the right. Scale bar 10 μm. **B**. Percentages of appropriately adhered and spread, uninfected and H37Rv infected M1 and M2 MDMs on crossbow micropatterns at 4, 8, 16 and 24 hours post infection. The number of cells that are adhered and spread in an appropriate manner per condition at 4 hours (h) is considered as the 100 %, with the following time points calculated as a percentage of the 4 hours. n=3 donors, mean ± SD.

In parallel to crossbow shaped micropatterns, we also made use of round shape micropatterns, to test whether the adherence and spreading rate depends on the shape of the micropatterns (Figure S3). Similar trends in all parameters were observed with round shaped micropatterns (Figure S3). Thus, we conclude that adherence does not depend on the shape of the micropattern when analyzing events up to 24 h post-infection, in both M1 or M2 macrophages. To summarize, we consider the H37Rv infection on MDMs on micropatterns assay to be best suitable for the study of early stages of H37Rv infection, up to at least 8 h to provide statistically robust measurements.

### Nucleus and MTOC positionings are altered in *Mtb* H37Rv infected M1 and M2 MDMs

One of the advantages of using micropatterns is that the position of organelles in adhered cells becomes typically maintained in a very constant manner, allowing reliable cell-to-cell comparison of organelle spatial relationships and their perturbance by a stimulus. In crossbow shaped micropatterned cells the nucleus is typically positioned in the center of the cell (Figure 4A) (Thery *et al*., 2006^a^). When artificially dividing the cell into four quadrants as shown in Figure 4A, the MTOC is typically found in the top polarized quadrant, creating an invisible axis from the nucleus, through the MTOC and towards the leading edge of the cell on the top of the crossbow. We sought to establish whether *Mtb* infection perturbs the typical nucleus and MTOC positioning, comparing H37Rv infected M1 and M2 MDMs to their respective uninfected control cells. To do so, we seeded M1 and M2 MDMs on crossbow shaped micropatterns, infected them with H37Rv and fixed 4 h post infection. We performed immunofluorescence with an anti-Tubulin antibody to visualize the microtubules and MTOC, labeled the DNA with DAPI and imaged the cells on a Zeiss LS880 airyscan confocal microscope, Zeiss Axio Observer 7, or StellarVision microscope. Typically, 60-70 % uninfected M1s and 65-75 % uninfected M2 in a sample display centralized positioning of the nucleus. We set the number of uninfected M1 and M2 MDM cells that displayed a centralized location of their nuclei as 100 % and compared the number of H37Rv infected M1 and M2 MDM cells to these 100 %, respectively.

**Figure 4.**
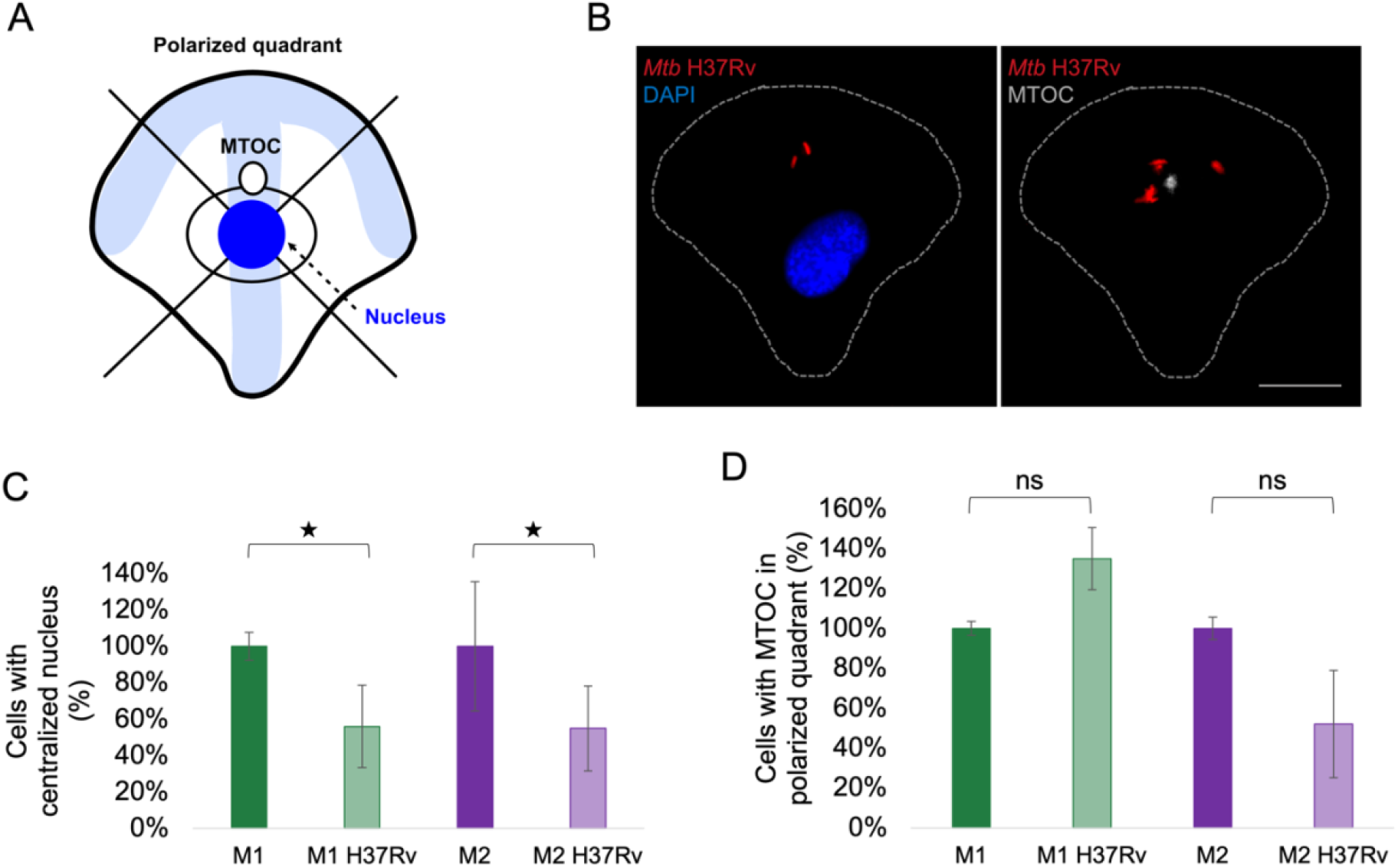
Nucleus and MTOC positionings are altered in H37Rv infected M1 and M2 MDMs. **A**. Schematic of a micropatterned cell with typical positions of the nucleus in blue and MTOC in white, as previously described (reference). Two artificial lines are drawn through the center of the cell to divide it into four quadrants. The top quadrant represents the polarized edge of the cell. **B**. Typical images of M1 MDMs on crossbow shaped micropatterns displaying correct nucleus and MTOC positions. DNA is labeled with DAPI (blue), the MTOC is marked in gray (microtubules and MTOC were stained with an anti-Tubulin antibody and the background microtubule staining was subtracted in FiJi to emphasize the MTOC), the contour of the cell is shown in a dotted line, *Mtb* H37Rv expresses mCherry (red). Scale bar 10 μm. **C**. The percentage of H37Rv infected MDMs relative to uninfected MDMs (set as 100 %), which display centralized location of the nucleus. **D**. The percentage of H37Rv infected MDMs relative to uninfected MDMs (set as 100 %), which display MTOC location in the polarized (top) quadrant. n=3 donors for M1 and n=2 for M2, mean ± SD. Analyzed by 2 tailed t-test; *, P < 0.05; ns, not significant

For both M1 and M2 H37Rv infected MDMs there was a notable reduction in the number of cells that displayed a typical, centralized position of the nucleus (56 % and 55 % for M1 and M2 MDMs, respectively, displayed centralized nuclear localization; M1 P=0.029, M2 P=0.023) (Figure 4C). In cells in which the nucleus was not found in the center, it was typically found off-center towards one of the right, left or top quadrants (see examples Figure S4B). Surprisingly, in M1 there was a 35% increase in the number of cells with the MTOC in the polarized quadrant when infected with H37Rv, relative to uninfected, whilst for M2 a 48% decrease in cells with the MTOC in the polarized quadrant was observed when H37Rv infected, relative to uninfected M2 (Figure 4D), with a trend towards significance compared to uninfected cells for M1 but not M2 (M1 P = 0.079, M2 P = 0.494). This suggests that even in H37Rv infected M1 cells in which the nucleus is not centered, the MTOC is more likely to be located in the correct quadrant, compared to infected M2. Taken together, this data suggests that infection with H37Rv might lead to alterations in positioning of cellular organelles both in M1 and M2 MDMs. Example images of centered nucleus and MTOC in H37Rv M1 and M2 MDMs are shown in Figure 4B and Figure S4A, respectively.

### Laboratory strain H37Rv and *Mtb* clinical isolate Ex52 localize in accordance with the MTOC and cellular polarity

Finally, we were interested to see whether once internalized, the *Mtb* bacilli were randomly distributed in space, or positioned in specific locations, which may be in line with the cellular polarity, measured by the nucleus-MTOC-cell edge axis. To do so, we infected crossbow micropatterned M1 and M2 MDMs from two donors with H37Rv and fixed samples 4 h post infection. We then performed immunofluorescence with an anti-Tubulin antibody to visualize microtubules and the MTOC, labeled the DNA with DAPI and imaged cells on a epifluorescence microscope. We manually measured the distance in pixels between each H37Rv bacilli and the MTOC, as described in the schematic in Figure 5A, with 40-225 H37Rv bacilli measured per condition (Figure 5B). Across two donors, the average distance of a H37Rv bacilli from the MTOC was significantly lower in M2s compared to M1s (mean distance 20.1 pixels versus 28.3 pixels, respectively in the first donor, P=3.9191 ×10^−12^, and 20.8 versus 27.8 pixels, respectively in the second donor, P=0.0053) (Figure 5B). The consistency in distance difference of bacilli from MTOC, between M1 and M2 suggests that there may be a functional consequence for the *Mtb* location relative to other organelles depending on cellular phenotype. Further experiments will need to be conducted to characterize the mechanistic insights of this phenomenon.

**Figure 5.**
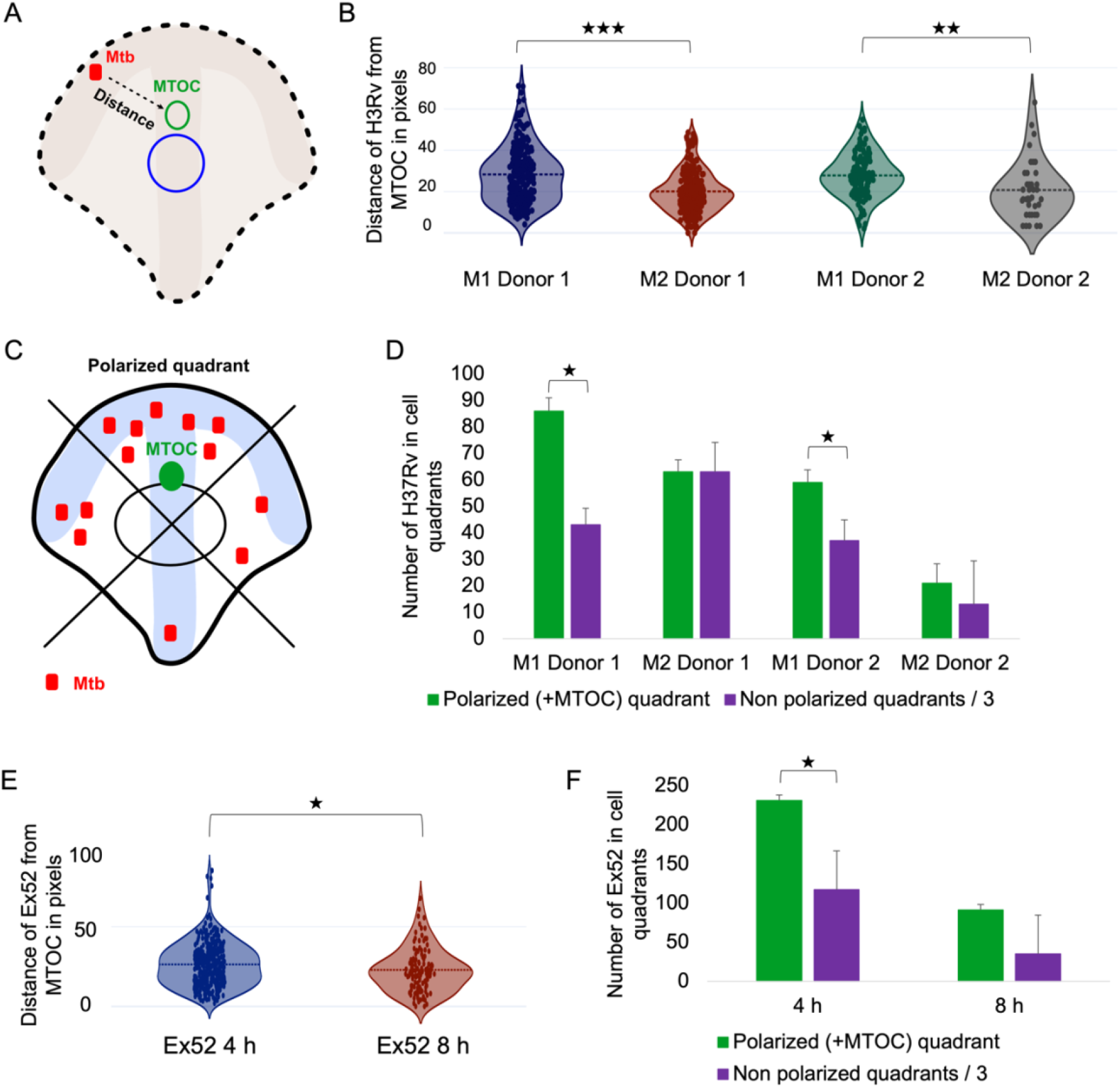
*Mtb* lineage 4 H37Rv and clinical isolate Ex52 localize in accordance to the MTOC and cellular polarity. **A**. Schematic describing the measurement of distance between each *Mtb* bacilli and the MTOC in pixels. **B**. Distance between H37Rv bacilli and the MTOC in pixels for two donors in M1 and M2 MDMs. **C**. Schematic describing distribution of *Mtb* bacilli in the different cellular quadrants. *Mtb* bacilli are counted in each quadrant and the average number of *Mtb* bacilli in a non-polarized quadrant is obtained by dividing the sum of all *Mtb* bacilli in the three non-polarized bacilli by three. **D**. The total number of H37Rv bacilli in the polarized quadrant compared to the average number of H37Rv in a non-polarized quadrant for M1 and M2 MDMs from two donors. **E**. Distance between the clinical strain Ex52 bacilli and the MTOC in pixels in M1 and M2 MDMs at 4 and 8 h post infection. **F**. The total number of Ex52 bacilli in the polarized quadrant compared to the average total number of EX52 in a non-polarized quadrant for M1 MDMs at 4 and 8 h post infection. Analyzed by 2 tailed T-test; *, P<0.05; **, P<0.01; ***, P<0.001, median plotted with SD.

Next, we counted the number of H37Rv bacilli in each quadrant (Figure 5C) to assess whether there is enrichment of H37Rv in the polarized quadrant. To do so, we divided the number of H37Rv bacilli in all three non-polarized quadrants by three, to account for the cell volume in each quadrant (assuming that the volumes of the four quadrants are roughly equal, see Savulescu *et al*., 2021) and compared this relative number to the number of *Mtb* H37Rv in the polarized quadrant. For M1 MDMs in both donors, a 2-fold enrichment of H37Rv bacilli was found in the total number of bacilli in the polarized quadrant compared to the total number of bacilli in non-polarized quadrants (P=0.046 and 0.020 for Donor 1 M1 and Donor 2 M1, respectively) (Figure 5D). Conversely, there was no *Mtb* enrichment in the polarized quadrant observed in M2 for either donor (P=0.207 and 0.216 for Donor 1 M2 and Donor 2 M2, respectively) (Figure 5D). Donor 2 M2 also had the lowest uptake of *Mtb* of all cells tested (21 versus 13 H37Rv in the polarized versus non-polarized quadrants). To summarize, this data suggests that coordinated spatial positioning of H37Rv relative to the MTOC and cell polarity differs between M1 and M2 MDM, as well as emphasizing donor variation that is inherent to human primary cell cultures and can be systematically quantified at subcellular levels using micropatterns.

Our data so far was conducted with the most commonly used lineage 4 laboratory strain *Mtb* H37Rv. We next sought to determine if the localization of *Mtb* bacilli in relation to the MTOC and cell polarity can also be observed, and remain consistent, with a lineage 4 clinical isolate of *Mtb*. For this, we seeded M1 MDMs on crossbow shaped micropatterns, infected them with clinical *Mtb* Ex52, which was transformed with a constitutive mCherry fluorescent reporter, and fixed them 4 h or 8 h post infection, to observe localization of Ex52 in relation to the MTOC and cell polarity over time. As for H37Rv, we measured the distance of each Ex52 bacilli from the MTOC, as well as counting the number of Ex52 in each quadrant. Interestingly, the mean distance of Ex52 bacilli from the MTOC was lower at 8 h versus 4 h post infection (20.8 pixels *vs*. 25.8 pixels, respectively, P= 0.0229) (Figure 5E), suggesting a potential tighter relative positioning with the MTOC and cell polarity over time. Lastly, we found the total number of Ex52 bacilli in the polarized quadrants to be almost 2-fold enriched comparing to the non polarized quadrants (231 *vs*. 117 bacilli, respectively at 4 h, P=0.016, and 91 *vs*. 35 bacilli, respectively, at 8 h, P=0.072), consistent with our observation with H37Rv in M1 at 4h (Figure 5F). Examples of Ex52 infected M1s in which the Ex52 bacilli are polarized in the polarized quadrant versus in the non-polarized quadrants are shown in Figure 5G. Taken together, this data confirms that for both the laboratory strain H37Rv and the clinical isolate Ex52, there appears to be a correlation with the MTOC positioning and cell polarity in M1 MDM.

## DISCUSSION

Here, we introduce a single cell-based approach to visualize and quantify host and pathogen factors which regulate *Mtb* phagocytosis by primary human macrophages and the spatiotemporal location of *Mtb* and host organelles in differential polarized macrophages, at a high resolution, in a high throughput manner. Our method is based on micropatterning of MDMs to reduce cell-to-cell variation, allowing for direct comparison of single cells. The use of micropatterns to study bacterial infection has been explored in the past, treating BMDMs with the edema toxin (ET) originating from *Bacillus anthracis* (Trescos *et al*., 2015) and infecting the HeLa cell line with *Salmonella Typhimurium* (*Stm*) (Vonaesch *et al*., 2013). However, to the best of our knowledge, this is the first time that bacterial infection using clinical and laboratory isolates is shown in micropatterned primary human macrophages.

We first sought to use elastomeric micropatterns assay to characterize a very early stage in *Mtb* infection of macrophages, namely phagocytosis. More specifically, we quantified the level of micropattern contraction as a measure of H37Rv phagocytosis. Indeed, cells infected with H37Rv displayed contraction, compared to uninfected cells, confirming that the elastomeric patterns assay can indeed be used to study phagocytosis of *Mtb* in MDMs. Furthermore, we showed that pre-treatment with IFN-γ increased contraction during *Mtb* infection, whilst conversely, decreasing expression of *NHLRC2* or removing surface lipids from *Mtb* by culturing in the presence of Tween-80, we can inhibit phagocytic contraction, and reduce bacterial uptake, measured by subcellular *Mtb* fluorescence. These findings exemplify a functional use for this assay to study the involvement of microenvironment modulation and host and pathogen factors in phagocytic processes on a single cell level and in a quantitative manner.

We then turned to standard micropatterns to study various aspects in *Mtb* infection. First, we optimized the assay to find the best suitable micropattern shape and size for MDM cells in general, and their performance for *Mtb* infected MDMs. We then characterized the time frame for infection using our assay. Our data showed that early, but not late stages of H37Rv infection can be studied in M1 and M2 MDMs, as cells start to detach from micropatterns at later time points. Interestingly, infection with H37Rv seemed to increase the adherence of M1, but not M2 MDMs on micropatterns. This observation is in line with previous studies that showed that *Mtb* infection increases cellular adhesion and surface presentation of adhesion molecules β2 integrin LFA-1 (CD11a) and its counter receptor ICAM-1 (CD54) in primary human macrophages (Rickenberg *et al*., 1975, DesJardin *et al*., 2002) and that M1 have higher expression of CD54, which also contributes to their rounder shape compared to elongated M2 (McWhorter *et al*., 2013).

As we propose our assay for the quantitative study of spatial subcellular localization of host-pathogen biomolecules and factors, we set out to explore whether infection of MDMs with the *Mtb* laboratory strain H37Rv affects the spatial positioning of key organelles. We specifically chose the nucleus of the cell, as it adopts typical positioning in micropatterned cells, as well as the MTOC, as it indicates the general cellular polarity. Interestingly, we observed opposite effects in the MTOC cellular positioning in M1 and M2 MDMs upon infection with H37Rv, namely an increase in the number of M1s that display typical MTOC positioning and a decrease in the number of M2s displaying typical positioning of the MTOC. These initial observations lay the foundation for future research to gain mechanistic insights into these potential differences and what impacts donor variability. One possible explanation could be the different level of heterogeneity between M1s and M2. More specifically, as M2s are more heterogeneous in shape, they would be expected to display more differences within a donor and between donors, depending on the level of differentiation of the cell. Typical centered location of the nucleus was observed in both M1s and M2 MDMs, however, in both cases, infection with H37Rv reduced the number of cells with centralized nuclei. This may suggest potential restructure of organelles or/and the cytoskeleton that might occur as a result of H37Rv infection, which can be investigated in future studies. Our findings demonstrate the utility of microfabricated patterns to investigate such phenomena and quantify the interaction between *Mtb* and organelle spatial organization.

In addition to studying the effects of H37Rv infection on organelle positioning, we also looked at the subcellular localization of H37Rv bacilli particularly in relation to the MTOC, which is indicative of cellular polarity. Interestingly, we found that the mean distance of H37Rv from the MTOC in M2 MDMs is shorter compared to M1 MDMs. We also found that there is an enrichment of H37Rv in the polarized quadrant of M1 MDMs compared to the non-polarized quadrants, but this was not consistently observed in M2 MDMs. Similar results for M1 were obtained with the clinical strain *Mtb* Ex52. Taken together, this data suggests that *Mtb* localizes internally in correlation to the cell polarity. It remains to be seen whether this preference for localization within the polarized cell space extends to its extracellular interactions, leading to preferred internalization of *Mtb* via the polarized edge, or whether the spatial organization occurs internally via a yet unknown mechanism. Imaging at earlier time points will address this question. Interestingly, although H37Rv is located further away from the MTOC in M1 MDMs compared to M2 MDMs, there is still a distinct enrichment of *Mtb* H37Rv in the polarized edge of the cell. A possible explanation would be that the early phagosome aligns with the nucleus-MTOC-leading edge axis and the general cellular polarization. Tran *et al*. offer another perspective in polarized epithelial cells infected with *Pseudomonas aeruginosa*. Bacterial aggregates on the apical membrane created a localized NF-κB host response and change in host spatio-temporal polarity, acting as an immune-activating signal. M1 MDMs may use a similar approach where *Mtb* is more enriched in the polarized quadrant to aid immune cell activation. Future research can gain further insights of this correlation between *Mtb* location and cellular polarity.

Taken together, by reducing cell-to-cell variability, we were able to compare between cells in a population and quantify subcellular localization patterns of *Mtb* in relation to the MTOC, as well as alterations in host organelle position post *Mtb* infection, and differences between individuals. These data would have been challenging to collect and quantify in standard cell culture, given the heterogeneity of the cell population. This is particularly the case with subtle changes in subcellular localizations of host and pathogen biomolecules or organelles, induced by *Mtb* infection. Furthermore, variability in qualitative and quantitative observations in host-pathogen interactions which occur between blood donors used for deriving primary macrophages are more readily assessed using micropatterning. The variability stems from variation in the genetic and epigenetic background of donors, as well as differences associated with donor gender, and age, which might affect host-pathogen interactions. Our protocol is particularly suited to study this variability in host pathogen interactions among the population, as by controlling the experimental system and reducing cell to cell variability, including cell size, shape, morphology and spatiotemporal localization of organelles, variability in host-pathogen interactions can be reduced to specific differences between donors.

To summarize, our assay is single-cell based and is suitable to study early stages of *Mtb* infection in MDMs in a quantitative and high throughput manner. Although our analysis was done manually, using FiJi, our assay can readily be coupled with automated analytical approaches, such as the one described in Savulescu *et al*., 2021. These analytical pipelines should be able to uncover patterns of subcellular distribution of biomolecules and organelles and enable quantitative, in depth characterization of spatial distribution patterns. This assay can be used to study the spatial distribution of a variety of pathogen or host derived biomolecules at early stages of *Mtb* infection of MDMs, in a quantitative manner, at the single cell level. Furthermore, we propose that this assay can be modified to study additional pathogenic infections in MDMs, or, alternatively, optimized to study *Mtb* infection in additional cell types.

## MATERIALS AND METHODS

### Human blood samples

Acquisition of human blood samples and the experiments that were performed with them were approved by the Human Research Ethics Committee of the University of Cape Town (HREC #317/2016). Buffy coats were obtained from anonymous blood donors through Western Province Blood Transfusion Service, Cape Town, South Africa.

### Generation of fluorescently-labeled *Mtb* and preparation of *Mtb* infection stocks

500 μg of pCherry3 vector (*smyc*’::mCherry; kind gift from David Russel (Tan *et al*., 2013) was electroporated into both H37Rv and clinical Ex52 and plated on 7H10/OADC (BD Biosciences) agar plates containing 50 μg/ml hygromycin. A single colony containing the fluorescently-labeled pCherry3 vector was inoculated into 10 ml 7H9/ADC (BD Biosciences) / 0.05 % Tween 80 (Sigma) media containing 50 μg/ml hygromycin and grown in static culture at 37 °C for 10 days. Cultures were frozen in aliquots in the presence of glycerol at -80 °C prior to the preparation of single cell stocks for H37Rv and clinical Ex52. A stock of H37Rv-GFP (a kind gift from the Molecular Mycobacteriology Research Unit, University of Cape Town, South Africa) was inoculated into 10 ml 7H9/ADC (BD Biosciences) / % Tween 80 (Sigma) media containing 25 μg/ml kanamycin and grown in static culture at 37 °C for 10 days. Cultures were also frozen in aliquots in the presence of glycerol at -80 °C prior to the preparation of single cell stocks.

For single cell stock preparation, frozen parent glycerol stocks were inoculated in 7H9/ADC (BD Biosciences) / 0.05 % Tween 80 (Sigma) media containing either 50 μg/ml hygromycin (H37Rv-pCherry3 and Ex52-pCherry3) or 25 μg/ml kanamycin (H37Rv-GFP) and grown in static culture at 37 °C for 10 days then sub-cultured 1/100 in the same medium in the presence of Tween80 (to generate H37Rv-Tween80) or absence of Tween 80 (to generate H37Rv and Ex52) for a further 10 days. Cultures were pelleted @ 3500 x *g* for 10 min, supernatant removed and pellets vigorously shaken with 2-3 mm glass beads for one minute. PBS was added to dispersed *Mtb*, which was gently pelleted @ 1400 x *g* for 10 min and the top half of the supernatant, containing single cell *Mtb*, was resuspended with a final volume of 5% glycerol and frozen at -80 °C. Before and after freezing, stocks were titrated and plated on 7H10/OADC (BD Biosciences) agar plates containing either 50 μg/ml hygromycin (H37Rv-pCherry3 and Ex52-pCherry3) or 25 μg/ml kanamycin (H37Rv-GFP) for CFU determination.

### Preparation of MDMs

Peripheral Blood Mononuclear cells (PBMCs) were obtained from healthy donor buffy coats, and were isolated using a Lymphoprep (Alere Technologies) density gradient. CD14 monocytes were isolated from PBMCs through CD14^+^ magnetic bead separation (MACS Miltenyi) and differentiated at 4×10^6^ cells per 40 mm dish for 7 days at 37 °C in RPMI media (Sigma) supplemented with 1 mM Sodium Pyruvate (Sigma), 2 mM L-Glutamine (Sigma), 10% Human AB Serum (hAB) (Sigma) and 5 ng/ml GM-CSF (MACS Miltenyi) or 100 ng/ml M-CSF (MACS Miltenyi), to generate the macrophage phenotypes M1 and M2, respectively. Following the 7 day differentiation, spent media was removed from plates and replenished with fresh prewarmed RPMI media (Sigma) supplemented with 1 mM Sodium Pyruvate (Sigma), 2 mM L-Glutamine (Sigma), 5 % hAB (Sigma) ± 100U/ml interferon gamma (PBL Assay Science) and incubated at 37 °C for 16-24 hours. Macrophages were then harvested by incubation with Accutase (Sigma) at 37 °C to gently detach cells followed by centrifugation at 300 x *g* for 10 minutes. Cell pellets were resuspended in fresh prewarmed RPMI media (Sigma) supplemented with 1 mM Sodium Pyruvate (Sigma), 2 mM L-Glutamine (Sigma), 5 % hAB (Sigma), counted and then diluted to a final concentration of ∼130 cells per microliter to prepare for cell seeding on micropatterns. For targeted gene-specific knockdown, MDM were seeded at 1×10^5^ cells/well and transfected with 60 nM total Silencer Select NHLRC2 (20 nM each of s51529, s51530 and s51531) siRNAs (Ambion) according to the method developed by Fisch *et al*, 2019, using the TransIT-X2 Dynamic Delivery System (Mirus Bio) for 48 hours followed by the addition of 100u/ml IFN-γ for 16-24 hours prior to harvesting and seeding MDMs onto micropatterns. As an siRNA control, 30nM scramble (4390843) siRNA (Ambion) was used.

### Preparation of microfabricated patterns (micropatterns)

Micropatterns were prepared based on the protocol described in Azioune *et al*., 2009. Shortly, we coated coverslips with PLL-g-PEG and then made use of a Cyto chromium photomask to imprint the desired size and shape of micropatterns onto coated coverslips. This was then followed by coating of the imprinted micropatterns with Fibronectin.

### Growing MDMs on micropatterns

Typically, 75,000 MDMs were seeded in a volume of 0.5 ml on coverslips containing fibronectin-coated micropatterns, in a 24-well plate. The cells were left to adhere for 1.5 hours, followed by removal of non adhesive cells by gentle washing with fresh prewarmed RPMI media (Sigma) supplemented with 1 mM Sodium Pyruvate (Sigma), 2 mM L-Glutamine (Sigma), 5 % hAB (Sigma), and infection with *Mtb* H37Rv at a multiplicity of infection (MOI) of 10 for 4 hours, or *Mtb* Ex52 at a MOI of 10 for 4 or 8 h. This was followed by fixation of cells for 16-24 hours with 4 % PFA, followed by 1x PBS washes and storage at 4 C for up to one week.

### Staining and immunofluorescence

Fixed samples were washed with 1x PBS, permeabilized for 10 min with 0.1 % Triton X-100, washed with 1x PBS and blocked with 3 % BSA in 1x PBS. Cells were then incubated with a primary rat monoclonal anti-tubulin antibody (Abcam, #ab6160) at 1:500 concentration for an hour, followed by three washes with 1x PBS and incubation with a secondary antibody conjugated to Alexa Fluor 647 (Abcam, #ab150151) at a concentration of 1:500. Samples were then washed with 1xPBS and incubated with μg/ml <u>DAPI</u> (4’,6-diamidino-2-phenylindole; Life Technologies, #D1306). Coverslips were then mounted in Vectashield (LSBio, #LS-J1032) and taken for imaging. For actin staining, cells were permeabilized for 10 min with 0.1 % Triton X-100, washed with 1x PBS and stained with Phalloidin–Atto 565 (SigmaAldrich, #94072). The coverslips were then washed with 1x PBS and incubated with μg/ml <u>DAPI</u> (4’,6-diamidino-2-phenylindole; Life Technologies, #D1306), followed by mounting as described above.

### Microscopy and image analysis

Mounted samples were imaged on a StellarVision microscope using Synthetic Aperture Optics technology, at a magnification of x20, on a Zeiss LSM880 airyscan confocal microscope, using a magnification of x60, and on a Zeiss Axio Observer 7 equipped with a Colibri 7 type RGB-UV LED illumination source, and controlled by ZEN 2.3 (Blue edition) software. For the data collected on the StellarVision, DAPI, AlexaFluor 488, mCherry and CY5 fluorescence were excited using 385/30 nm, 469/38 nm, 555/30 nm and 631/33 nm, respectively. Raw 16 bit images were saved as .tif files prior to image processing and analysis. For the data collected on the Zeiss Axio Observer, bright-field and fluorescent images were captured using a Zeiss Axiocam 506, using a Plan-Apochromat 100X (1.4NA) Phase Contrast Objective. DAPI, AlexaFluor 488, mCherry and CY5 fluorescence were excited using 385/30 nm, 469/38 nm, 555/30 nm and 631/33 nm, respectively. Raw 16 bit images were saved as .CZI files prior to image processing and analysis. For the data collected on the Zeiss LSM880 airyscan microscope, DAPI, AlexaFluor 488, mCherry and CY5 fluorescence were excited using 385/30 nm, 469/38 nm, 555/30 nm and 631/33 nm, respectively. All images were processed and analyzed with FiJi. Image processing included background subtraction and analysis included measurement of distance between MTOC and Mtb bacilli, superimposing the quadrant division and quantification of Mtb bacilli per quadrant and measurement of fluorescence intensity.

## ACKNOWLEDGMENTS

We would like to thank Dr. Caron Jacobs and the UCT Infectious Disease and Molecular Medicine Microscopy Unit, as well as the SAMRC/NHLS/UCT Molecular Mycobacteriology Research Unit for the use of their microscope for this work. We would also like to thank Prof. Edward Sturrock for providing lab space and for continuous support.

## COMPETING INTERESTS

The authors declare no competing interests.

## AUTHOR CONTRIBUTIONS

Conceptualization: A.F.S., A.K.C., M.M.M.; Methodology: A.F.S., D.O., N.P., R.H.; Validation: A.F.S., N.P.; Formal analysis: A.F.S.; Investigation: A.F.S., N.P.; Writing - original draft: A.F.S.; Writing - review & editing: A.F.S., D.O., N.P., A.K.C., M.M.M; Supervision: A.K.C., M.M.M.; Project administration: A.F.S., A.K.C., M.M.M., N.P.

## FUNDING

This research has been supported by the following grants: A grant from the Emerging Research Area Program of The Department of Science and Technology (DST, South Africa) Department of Science & Technology Centre of Competence Grant, SA Medical Research Council SHIP grant, WUN CIDRI Ph.D. scholarship and the Carnegie Corporation DEAL fellowship.

## SUPPLEMENTARY FIGURES

**Figure S1.**
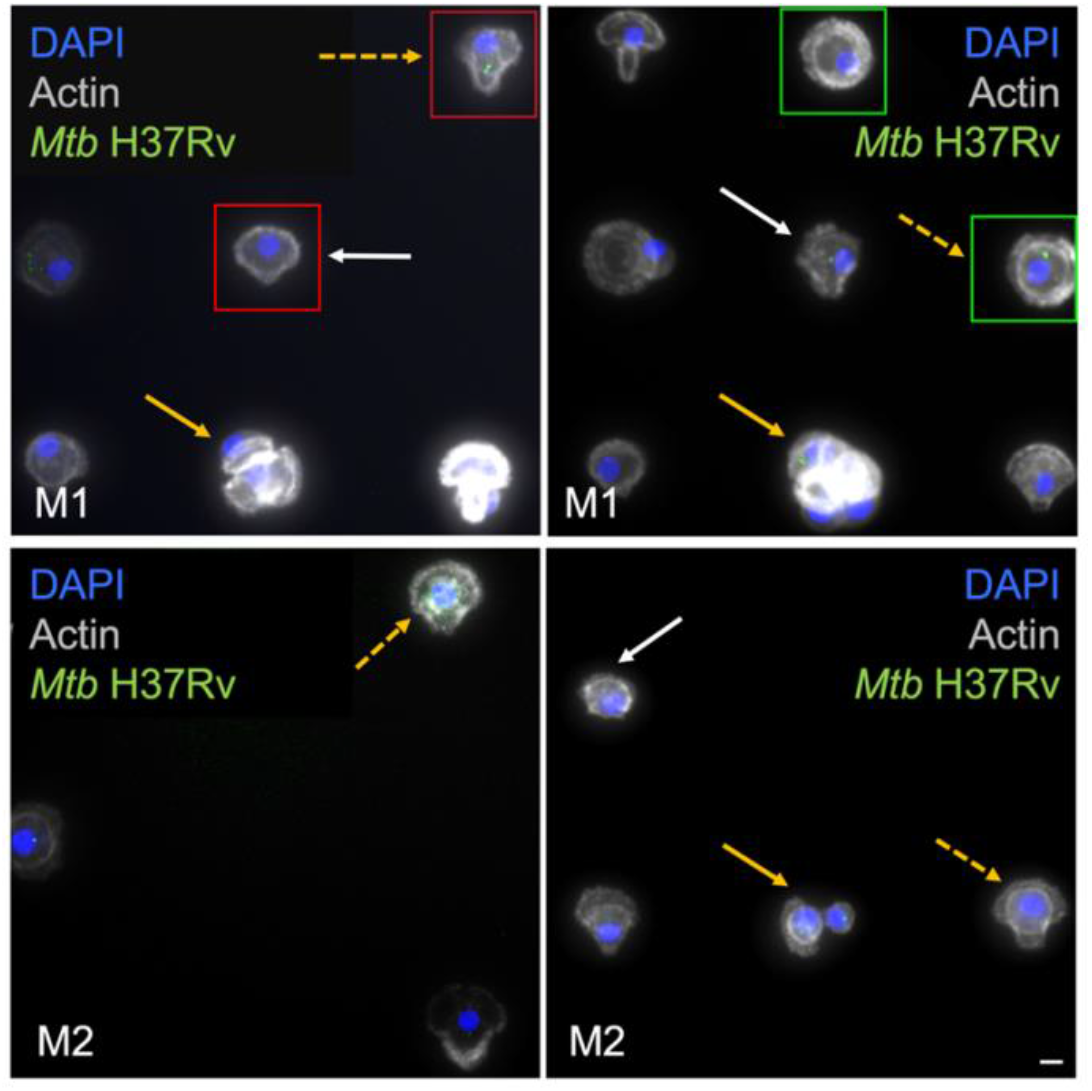
Optimization of the protocol on round versus crossbow shaped micropatterns. M1 and M2 MDM cells were seeded and grown on medium sized crossbow (examples in red squares) or round shaped (examples in green squares) micropatterns and infected with H37Rv, as detailed in Figure 1A. Orange dotted arrows indicate cells that spread properly on micropatterns, orange arrows indicate multiple cells on one micropattern and white arrows indicate cells that did not spread properly on micropatterns. The DNA is stained with DAPI (blue), H37Rv is tagged with GFP (green) and actin is stained with phalloidin (gray). Scale bar 10 μm.

**Figure S2.**
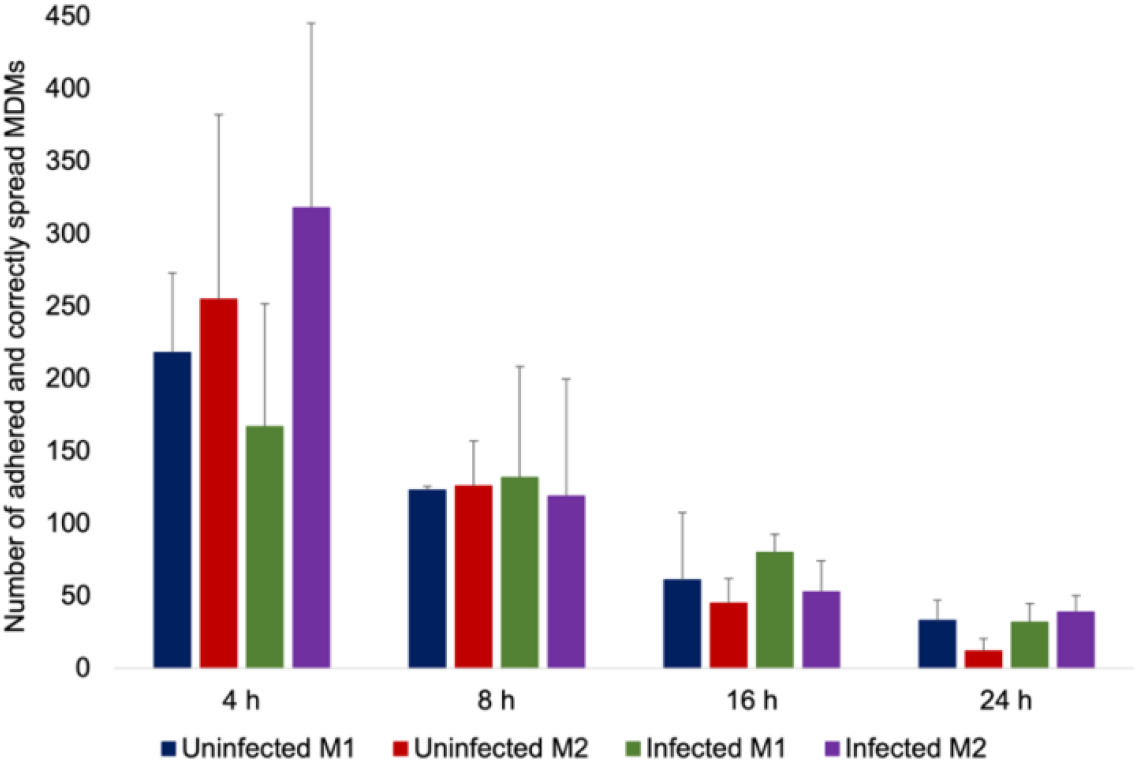
Absolute number of MDMs adhered and correctly spread on crossbow shaped micropatterns at different time points post infection with H37Rv. n=3 donors, mean ± SD. P=0.832 for uninfected M1s vs M2s at 4 h; P=0.476 for infected M1s and M2s at 4 h; P=0.725 for uninfected versus infected M1s at 4 h; P=0.772 for uninfected versus infected M2s at 4 h. Analyzed by 2 tailed T-test.

**Figure S3.**
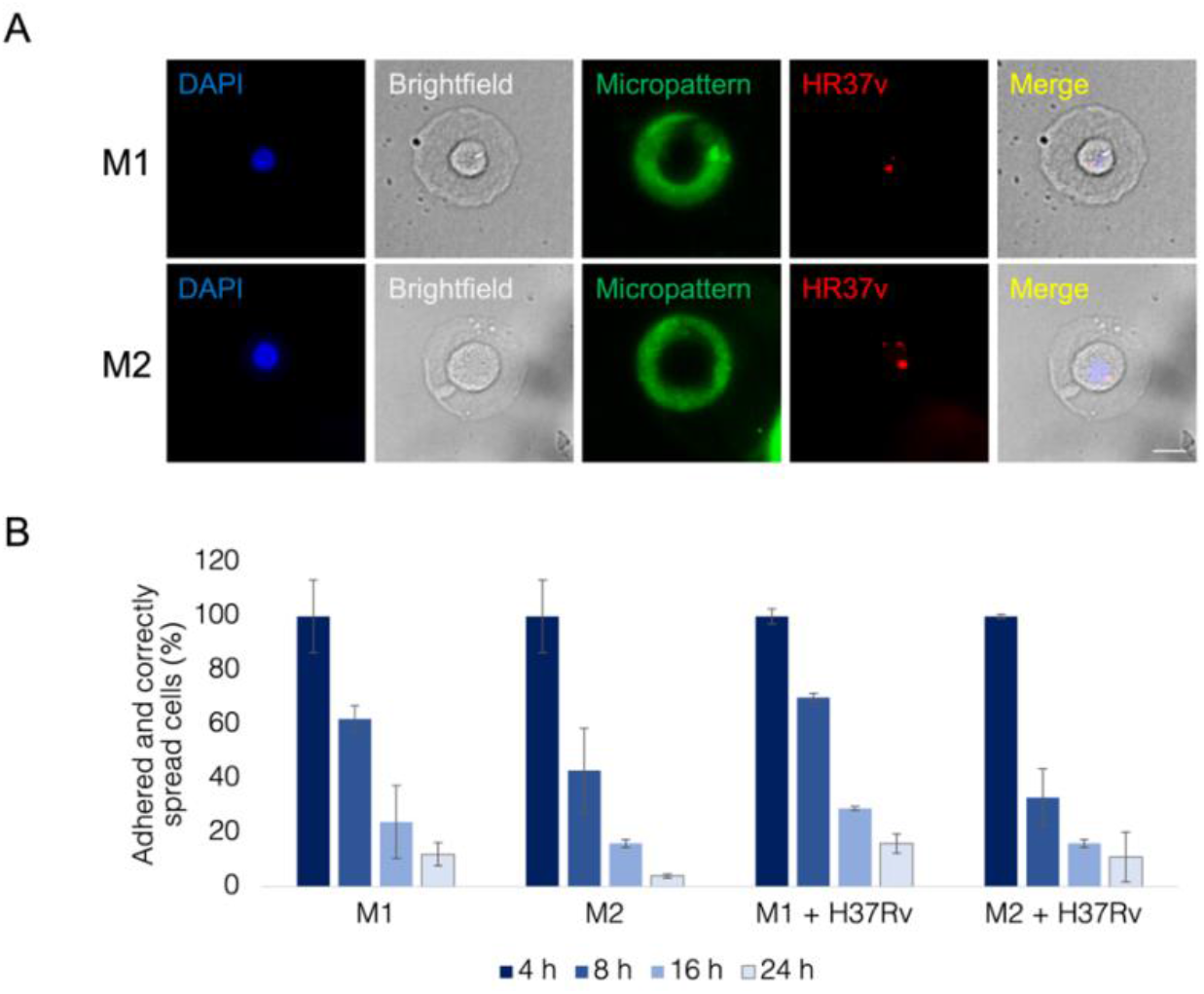
Percentages of appropriately adhered and spread, uninfected and H37Rv infected M1 and M2 MDMs on round shaped micropatterns at 4, 8, 16 and 24 hours post infection. The number of cells that are adhered and spread in an appropriate manner per condition at 4 h is considered as the 100 %, with the following time points calculated as a percentage of the 4 hours. n=3 donors, mean ± SD.

**Figure S4.**
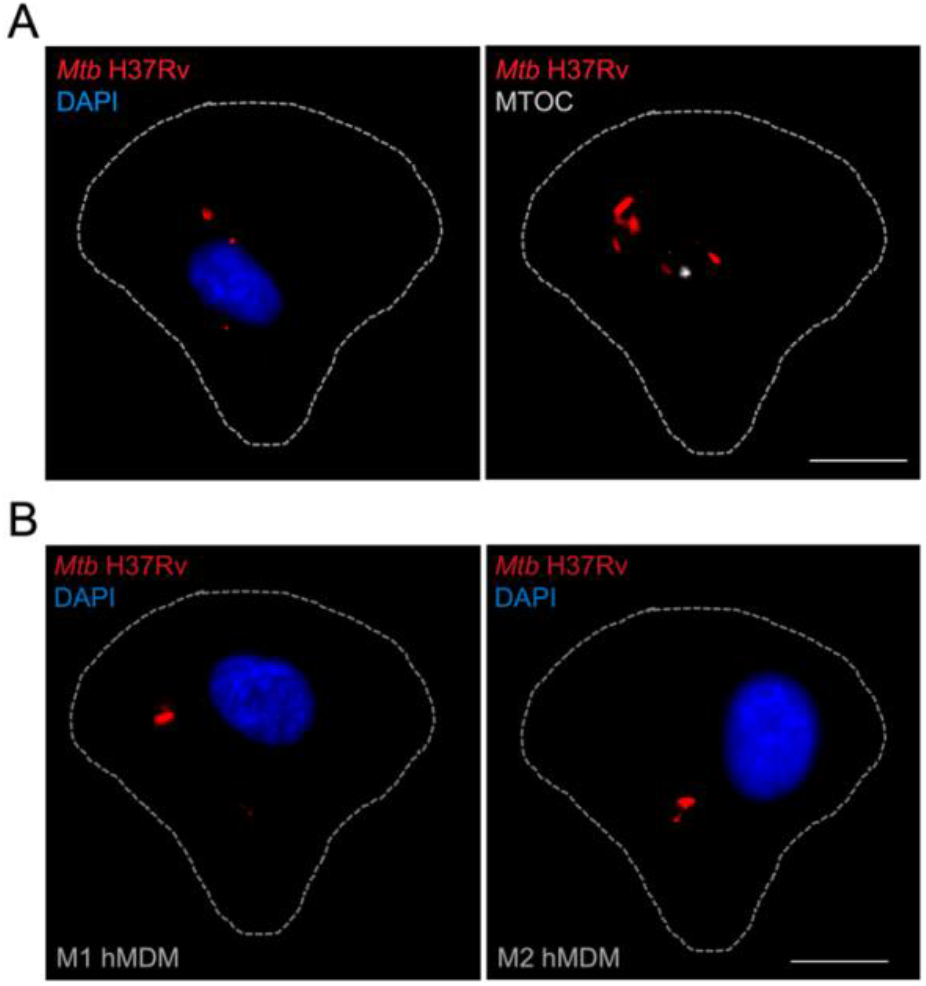
Example images of H37Rv infection in MDM. **A**. M2 MDM cells infected with H37Rv where the nucleus (left image) and MTOC (right image) are positioned correctly. The DNA is stained with DAPI (blue), H37Rv is tagged with mCherry (red) and the MTOC is marked in gray (microtubules and MTOC were stained with anti Tubulin and the background was removed to emphasize the MTOC), the contour of the cell is shown in dotted line. Scale bar 10 μm. **B**. Example images of H37Rv infected MDMs (M1 on the left and M2 on the right) in which the nucleus is not positioned correctly. The DNA is stained with DAPI (blue), H37Rv is tagged with mCherry (red), scale bar 10 μm.

## Notes

### Competing Interest Statement

The authors have declared no competing interest.

